# Citizen science data indicates morphological complexity of galls depends on the originating plant organ

**DOI:** 10.1101/2025.05.17.654643

**Authors:** Kanako Bessho-Uehara, Riki Takara, Kosuke Sano, Kohei Tamura

## Abstract

Galls are abnormal plant structures formed through interactions between host plants and insects, providing shelter and nutrients for gall-inducing insects. As distinct insect species can generate unique gall morphologies even on the same host plant, galls are often viewed as an extended phenotype of the insect. However, since galls consist of plant-derived cells, plant factors are also hypothesized to shape their morphology. Previous studies exploring this possibility have been restricted to one or a few plant species, limiting broad evolutionary inference. Here, we used citizen science observations to analyze gall morphological complexity across 26 plant orders. Quantitative comparisons using fractal dimension indices revealed that stem-derived galls display significantly less morphological variation than leaf-derived galls. Generalized linear mixed models indicated that stems possess lower morphological plasticity than leaves. These results held even after accounting for insect and plant phylogeny, suggesting that gall form is influenced by both insect species and the developmental properties of the host organ. Our findings highlight the role of plant organ identity in modulating gall morphology and demonstrate that tissue plasticity constrains insect-induced developmental outcomes. This study provides the large-scale cross-species analysis of gall formation and illustrates the power of citizen science in studying morphological evolution across taxa.

## Introduction

Citizen science has emerged as a powerful tool for biodiversity and ecological studies^1^. Online platforms enable the public to share observations of organisms, generating a vast amount of biodiversity data that significantly expands the spatial and temporal scope of scientific research^2^. An influx of publicly contributed data has revolutionized multiple research fields, allowing scientists to reveal large-scale species distribution patterns at unprecedented scales^3^. One such tool is iNaturalist (https://www.inaturalist.org/), a community-driven, free-access platform that allows users to share observations of animals, plants, and other organisms through a dedicated website and mobile application^4^. Users can upload photos of organisms to iNaturalist, and AI-powered identification suggests possible species, which are then verified by a community of experts and enthusiasts. This approach enables quick and accurate species identification, and iNaturalist has gained increasing recognition among researchers worldwide for its potential applications in biodiversity and conservation researches^5^. Recent studies have integrated iNaturalist data for species distribution modeling and biodiversity assessment across various taxa. For example, how to track the spread of exotic species and examine geographic variation in traits like color changes in rat snakes depends on the latitude^6^, melanism in monarch butterfly larvae^7^, and roadkill patterns in rattlesnakes^8^. Furthermore, iNaturalist has facilitated documentation of rare species^9,10^ and even the discovery of new ones^11^. Whereas iNaturalist enables instantaneous collection of large-scale global data, community science data must meet quality standards to be effectively utilized in research, as observer bias and data variability can affect the results^12,13^. To reduce these concerns, advanced statistical approaches have enabled researchers to extract meaningful conclusions from citizen science datasets. This methodology has proven to be particularly effective in biogeography, species monitoring, and conservation studies^14–16^. Yet, its application to morphological research and species interactions remains relatively underexplored.

To investigate the potential of citizen science data in morphological studies, we focused on insect-induced plant galls—specialized structures formed by complex plant-insect interactions^17,18^ (**Fig. 1a**). Gall formation is a dynamic and highly specialized process, driven by a combination of mechanical stimuli, chemical stimuli, and genetic responses of insects and plants^19^. Gall morphology serves important ecological functions for gall-inducing insects, including regulating the microenvironment, nutrient acquisition, and defense against natural enemies^20,21^. Traditionally, gall morphology has been attributed to the inducing insect species, as different insects generate distinct gall structures on the same host plant^22,23^. That is why the galls are considered the `extended phenotype` of gall-inducing insects^21^. Although gall-inducing insects play a pivotal role in shaping these structures, the developmental properties of plant tissue from which they arise are likely to influence their final morphology. Previous studies have shown that closely related insect species can induce morphologically diverse galls depending on the plant organ they affect^24^. A different study suggested that stems have lower morphological plasticity than leaves, potentially resulting in more constrained morphologies of galls^25^. Such observations suggest that morphological diversity of galls varies based on the originating plant organ, which may impose structural constraints on gall development, offering a different perspective from the conventional views. However, such studies have been limited to single or few plant species, leaving a gap in our understanding of whether this pattern holds across diverse plant taxa.

**Figure 1.**
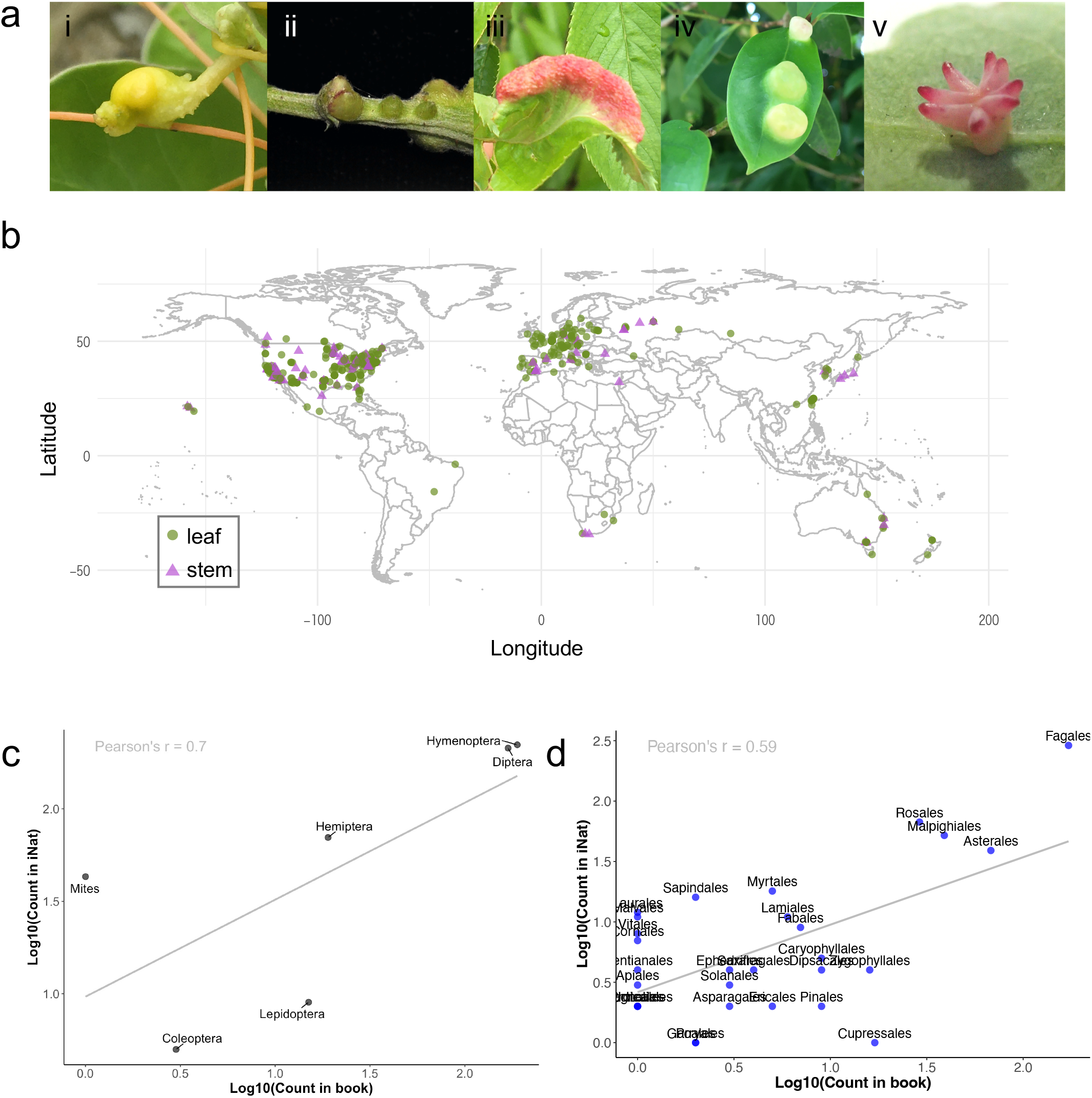
Information about gall images retrieved from iNaturalist. (a) Pictures of several galls originated from the stem and leaf. The species name of plant – insect pairs as i, *Cuscuta campestris – Smicronyx madaranus*; ii, *Artemisia princeps – Rhopalomyia struma*; iii, *Prunus ×yedoensis – Tuberocephalus sasakii*; iv, *Distylium racemosum – Neothoracaphis yanonis*; v, *Quercus lobata – Cynips douglasii* (b) Geographic distribution of the acquired observations on iNaturalist. (c-d) The scatter plot illustrates the relationship between the log-transformed counts of insect (c) or plant (d) orders recorded in book data and iNaturalist data. Each point represents an order, with its name displayed next to the corresponding point. The solid gray line represents the linear regression trend, and the Pearson correlation coefficient is displayed in the upper-left corner.

To address this limitation, we leveraged the iNaturalist data to conduct a large-scale comparative analysis of gall morphology. By analyzing hundreds of gall observations spanning 26 plant orders, we quantitatively assessed morphological complexity using fractal dimension analysis and tested whether originating plant organ imposes constraints on gall development. Additionally, we applied Markov chain Monte Carlo generalized linear mixed models (MCMCglmm) to evaluate the relative influence of plant organ on gall shape while accounting for phylogenetic relationships. These results offer support for the notion that gall morphology is influenced not only by the inducing insect but also by structural constraints imposed by the host plant organ—an aspect that has received less attention in previous studies. In addition, by incorporating large-scale citizen science data, this study demonstrates the potential of community-sourced biodiversity observations for addressing morphological questions across diverse taxa.

## Results

### iNaturalist-derived data show no strong sampling bias

We first retrieved observation records tagged with the keyword “gall” from iNaturalist (**Fig. S1**). Most records originated from North America and Europe, although some originated from Brazil, Asia, and Australia (**Fig. 1b**). Because iNaturalist is an unstructured, user-driven platform, its data are susceptible to observer bias^26^. On the contrary, the illustrated book represents structured sources and often offer more comprehensive coverage of local taxa. To evaluate the randomness of data sampling, the proportions of host plants and gall-inducing insects between from iNaturalist and the gall book. As most of the iNaturalist-derived data were from North American and European sources, the comparison was made to with taxa published in North America (‘Galls of Western United States’)^27^. We compared the proportions of each insect and plant order, divided into stem-derived and leaf-derived galls. As for plant orders in total, 26 orders were included in the iNaturalist-derived data (**Fig. S2a**) and 22 orders were included in those derived from the gall book (**Fig. S2b**). This difference in numbers is probably due to the difference in the size of the area of the acquired data. The number of insect orders retrieved from both data sources were same, six orders (**Fig. S2c, d**). Diptera were the most common in iNaturalist stem galls (64.2%), while Hymenoptera dominated in the book-derived data (44.8%). Despite these differences, the overall correlation in taxonomic frequencies was high (*r = 0*.*97* for insects, *r = 0*.*95* for plants; **Fig. S2e, f**). Log-transformed counts also showed strong correlation (*r = 0*.*70* for insect order, *r = 0*.*59* for plant order) (**Fig. 1c, d**), suggesting minimal sampling bias in the iNaturalist dataset.

### Diversity of combinations between gall-inducing insects and host plants

Galls are formed by the combinations of insect and plant species. The number of combinations of insect and plant species at the order level is suggested that most combinations of insect and plant species are less than five (**Fig. 2a**). A heat map presenting the relationship between gall-inducing insect and host plant phylogeny showed that Hymenoptera–Fagales was the most frequent pairing, followed by Diptera–Fagales (**Fig. 2b**). This result is consistent with reports that the insect galls induced by the Cynipidae and Cecidomyiidae are found on many Fagaceae plants^28^. Diptera showed the broadest host range, while Lepidoptera was found to be parasitic on a specific group of plant species (Caryophyllales and Asterales) (**Fig. 2b**). To clarify whether these specific insect–plant combinations determine the diversity of gall morphologies, fractal dimension analysis was performed. Fractal dimension provides a quantitative, scale-independent measure of morphological complexity, capturing structural irregularity and self-similarity more effectively than traditional shape descriptors^29–31^. This metric allows for objective comparisons across different sizes and biological forms, making it particularly suitable for analyzing intricate structures such as leaves and root morphologies^32,33^. We quantified the fractal dimension values of all galls and summarized the results by insect order. Values ranged from 1.0 to 1.2 across all orders, with Diptera reaching up to 1.4, indicating its potential to induce highly complex galls (**Fig. S3a**). However, no significant differences in fractal dimension were detected among insect orders (Wilcoxon rank-sum test, *p > 0*.*1*). To further examine whether particular insect–plant combinations generate distinct morphological patterns, we visualized the mean fractal dimension for each insect–plant order pair (**Fig. S3b**). While some combinations— such as Diptera–Fagales or Mites–Myrtales —appear to yield higher or lower complexity on average, the overall pattern does not indicate any consistent or systematic relationship. That is, morphological complexity varied across combinations but without clear phylogenetic structuring or interaction patterns. This suggests that, although individual cases of high or low complexity exist, the average gall morphology might be difficult to determine by the specific combination of insect and host plant lineages.

**Figure 2.**
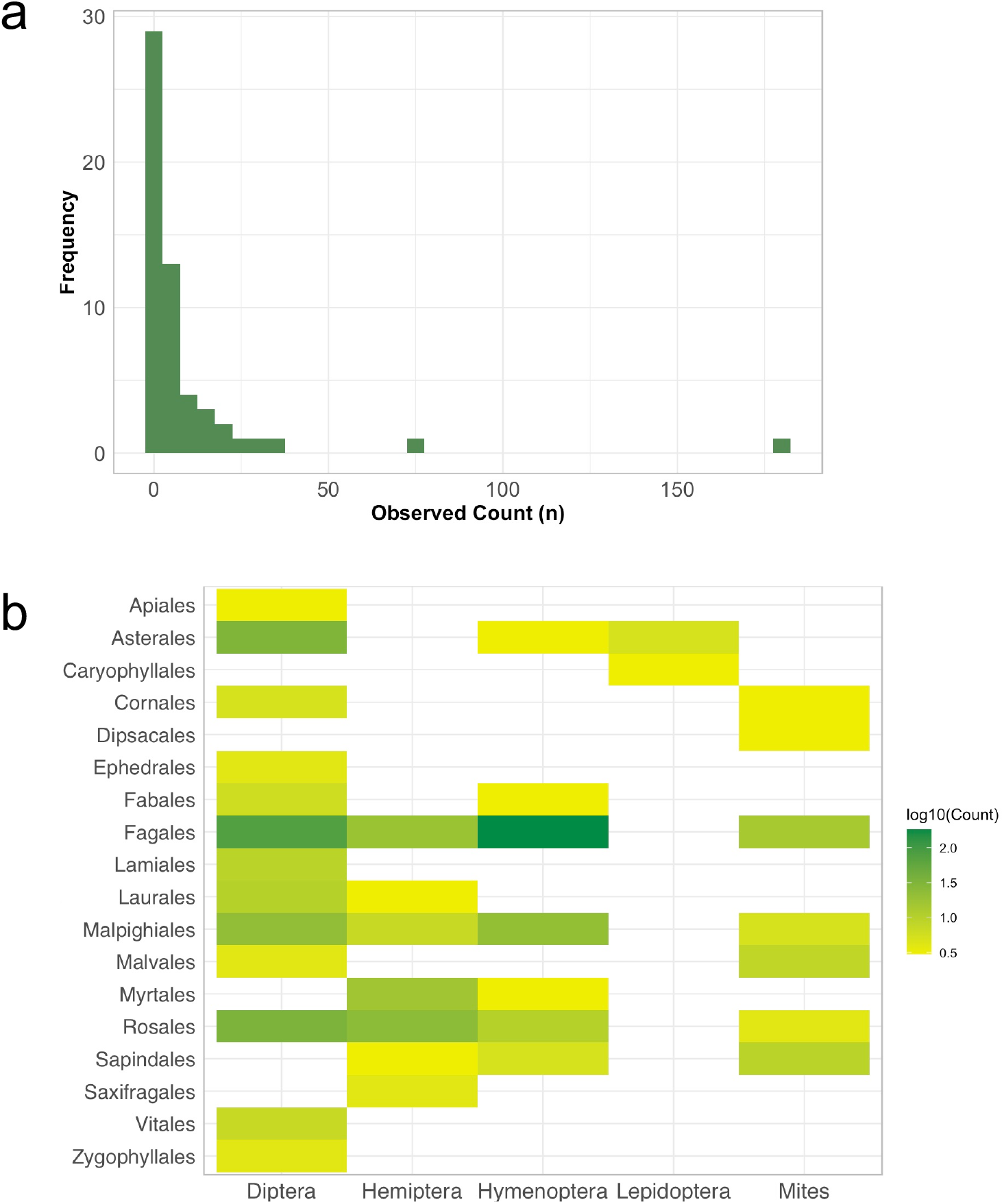
Relationships between gall-inducing insects and host plants. (a) Distribution of the number of combinations between gall-inducing insects and host plants. (b) Heatmap showing the number of observed combinations between gall-inducing insect and plant orders. The color scale represents the base-10 logarithm of the number of observed insect–plant interactions per order combination. Combinations with only a single observation (n = 1) were excluded to minimize sampling noise.

### Stem-derived galls have less morphological diversity than leaf-derived galls

Given that morphological variation was not significantly different among insect orders, we next examined whether the plant organ of gall origin influences morphological diversity. Galls were categorized as leaf-derived or stem-derived, and their fractal dimensions were compared. The images of stem-derived galls were taken from the side. However, many images of leaf-derived galls were taken not only from the side but also from above or diagonal angle. In order to compare differences in morphological diversity depending on the photo angle, images of leaf-derived galls taken from three different angles were used. Fractal dimension values ranged from 1.0 to 1.4 for leaf-derived galls and from 1.0 to 1.2 for stem-derived galls. Median values were 1.08 for leaf-derived galls (regardless of angle) and 1.06 for stem-derived galls (**Fig. 3**). This variation in morphological diversity was significantly greater for leaf-derived galls, indicating that leaf-derived galls are more complex compared to those derived from stems (Wilcoxon rank sum test, *p < 0*.*05*). To test whether this pattern holds within insect groups, we analyzed galls induced by Diptera and Hymenoptera—the two most abundant taxa in our dataset. In both groups, leaf-derived galls showed significantly greater complexity than stem-derived galls (**Fig. S4a, 4b**), suggesting that the influence of plant organ on gall morphology persists regardless of insect identity.

**Figure 3.**
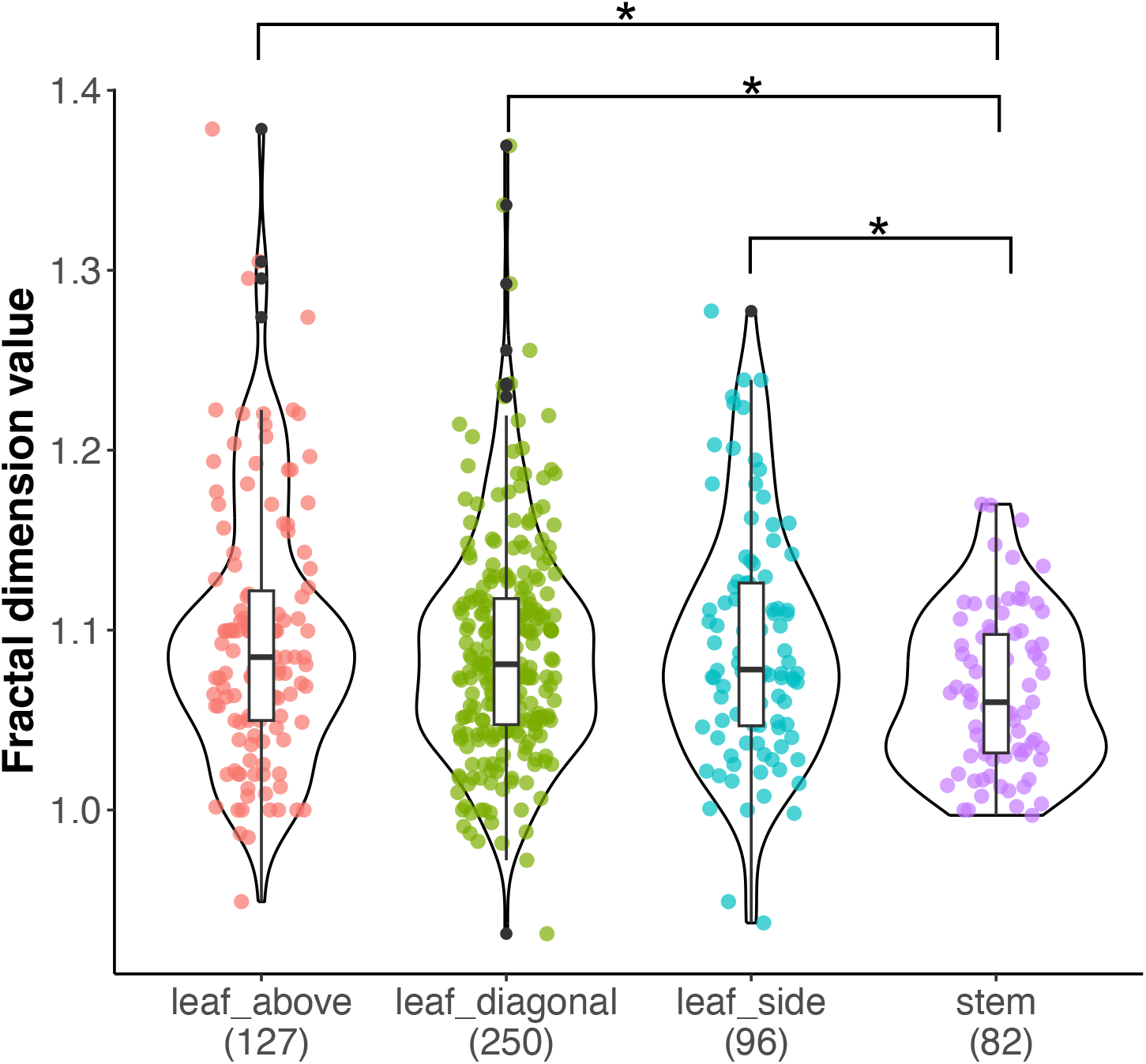
Fractal dimension analysis of gall morphological diversity depends on the originating plant organ. Pairwise comparisons using Wilcoxon rank sum test with Bonferroni correction. *p-value <0.05.

### MCMCglmm revealed that stem has less plasticity compared to the leaf

Having found that the originating plant organ influences gall morphological variation, we further quantified the effect of plant organ on gall morphology using a phylogenetically informed Bayesian mixed model (MCMCglmm). The response variable was the fractal dimension of gall morphology, and the explanatory variable was plant organ (leaf vs. stem). To account for potential phylogenetic non-independence, we incorporated phylogenetic relationships among host plant orders and gall-inducing insect orders as random effects. This modeling framework allowed us to estimate the fixed effect of plant organ while statistically controlling for shared evolutionary history (i.e., phylogenetic inertia). The MCMCglmm analysis revealed that stem-derived galls exhibited consistently lower fractal dimension values than leaf-derived galls, even after correcting for phylogenetic structure (**Fig. 4, Table S3**). The effect of organ type was statistically significant (p_MCMC_ < 0.001), with the posterior distribution of the stem effect clearly separated from zero, indicating a robust negative effect of stem-derived origin on gall complexity. Furthermore, the contributions of host plant and insect phylogenetic structure, modeled as random effects, were smaller than the organ effect, highlighting the role of developmental constraints imposed by plant organ in shaping gall morphology across diverse taxa.

**Figure 4.**
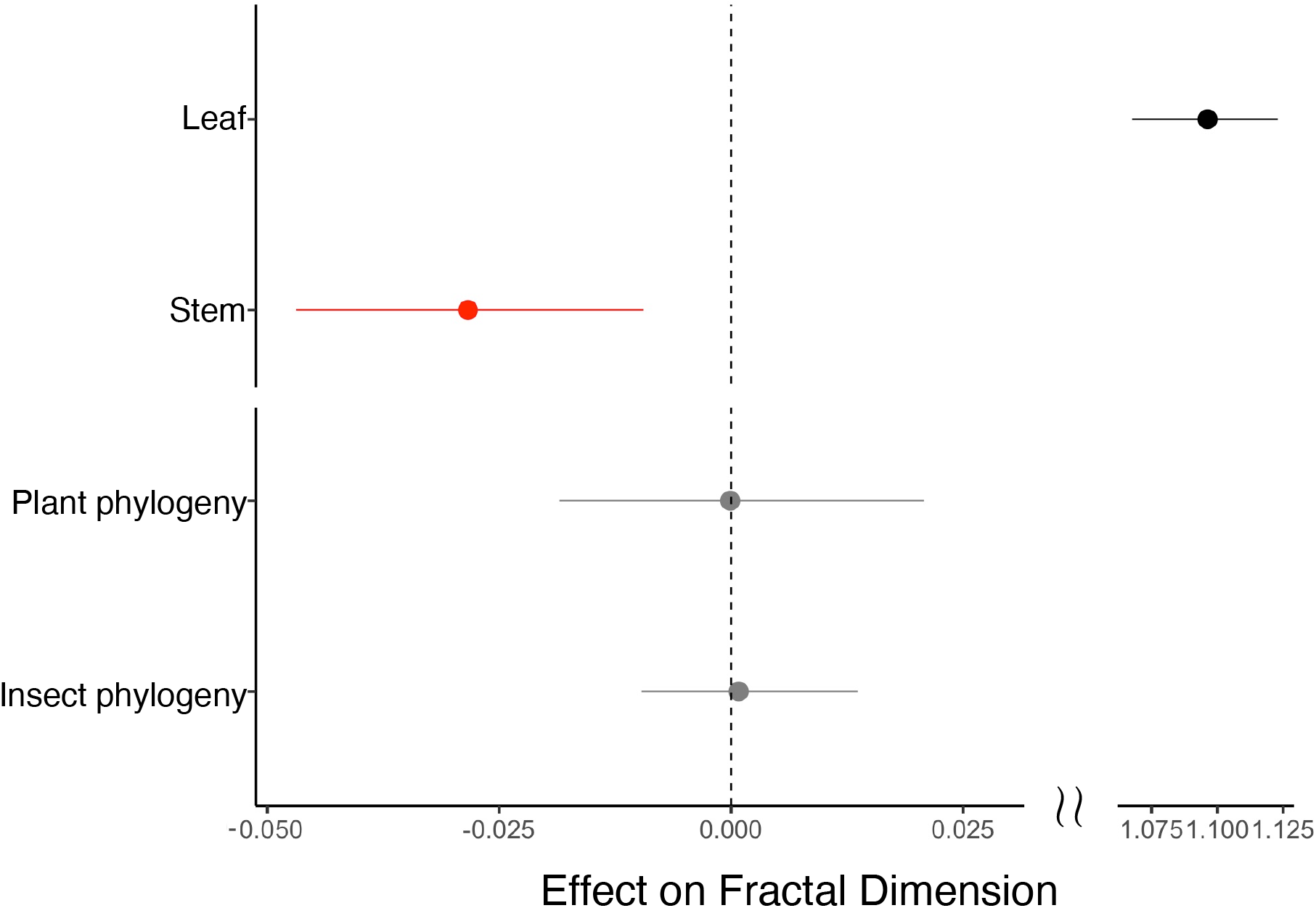
Coefficients of originating plant organs for gall morphological diversity. The horizontal axis represents the coefficients obtained from the MCMCglmm analysis, showing the fixed effects (originating plant organ; leaf and stem) and random effects (phylogeny of plant and insect). They indicate the 95% credible intervals, while the points represent the mean values. An asterisk denotes significant coefficients, where the 95% credible interval does not overlap with zero.

### PCA reveals contrasting morphological traits between leaf- and stem-derived galls

While fractal dimension analysis was useful for quantifying overall morphological complexity across structurally diverse galls, it required analysis based on binarized images, which inevitably resulted in the loss of certain morphological information. To complement this limitation and assess whether image-based features align with human-perceived complexity, we quantified six morphological traits and conducted principal component analysis (PCA). The traits measured were: (1) aspect ratio, (2) thickness of the gall relative to its organ of origin (relative thickness), (3) deviation from an ideal ellipse (variance from ellipse), (4) number of thorns, (5) color difference from the source organ, and (6) trichome density and length (Fig. 5a). For this analysis, only side-view images of galls were used to ensure comparability in assessing aspect ratio and thickness. The PCA plot revealed a clear separation between stem- and leaf-derived galls, primarily along PC1 (Fig. 5b). Stem-derived galls clustered mainly on the negative side of PC1, whereas leaf-derived galls were more widely distributed across the PCA. Leaf-derived galls were more strongly associated with qualitative traits such as color change and trichome density, as well as quantitative features including relative thickness and deviation from elliptical shape. These findings suggest that plant organ influences gall morphology in terms of both qualitative and quantitative traits.

**Figure 5.**
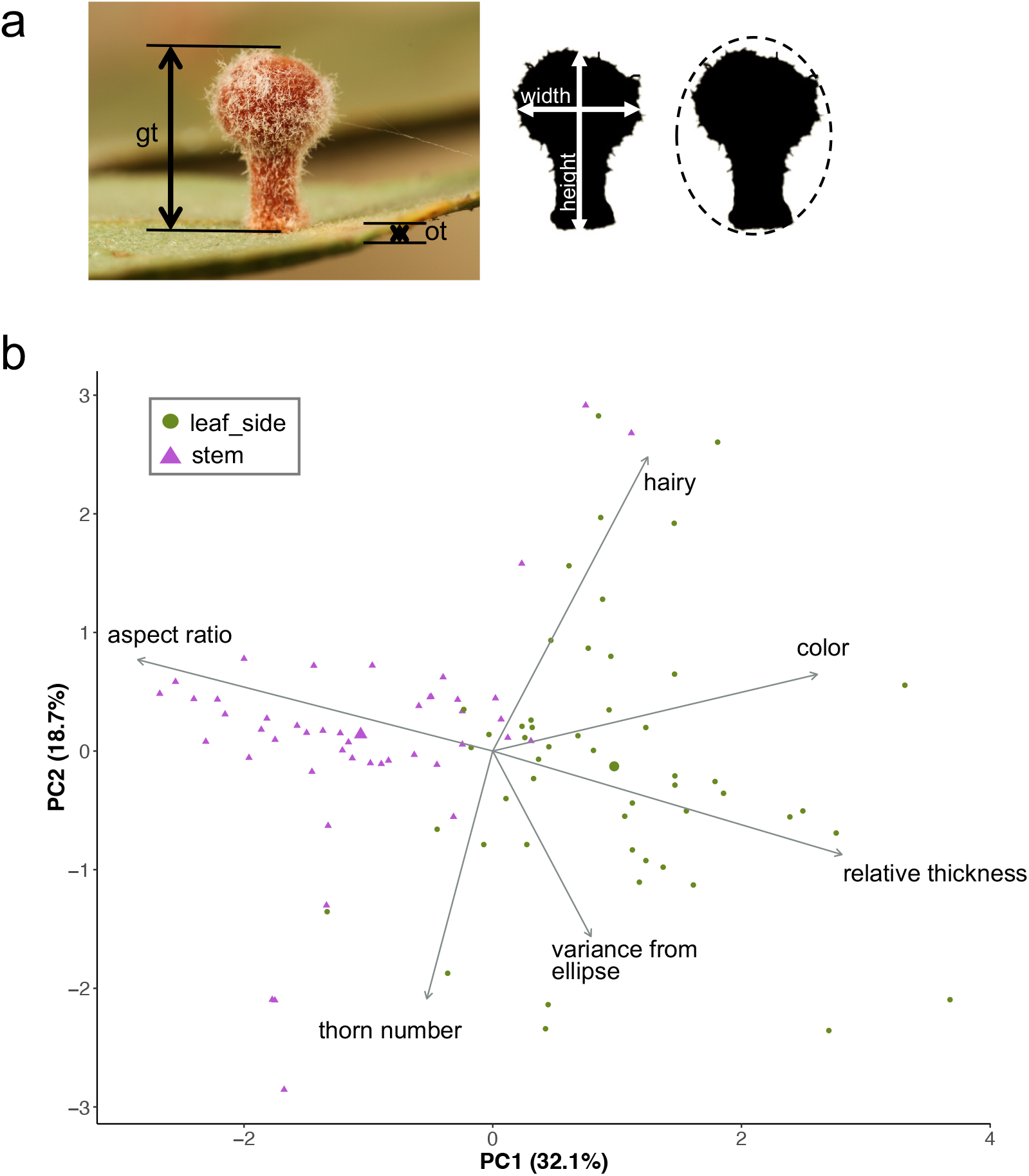
Principal component analysis using several characteristics of galls. (a) Diagrams on how to measure the factors for principal component analysis. Gall image was retrieved from iNaturalist (https://www.inaturalist.org/observations/8129878). The diagrams showed how to calculate relative thickness, aspect ratio, and variance from the ellipse. Gt, gall thickness; ot, organ thickness (in this case, the thickness of the leaf blade). (b) Biplot of the result of PCA visualizing the variance and distribution of samples based on originating plant organs. Arrows indicate PCA loadings.

## Discussion

This study provides evidence that the plant organ influences gall morphological complexity, with leaf-derived galls exhibiting greater diversity compared to stem-derived galls. Our results support the hypothesis that plant tissue plasticity plays a key role in shaping gall morphology, expanding to the traditionally insect-centric view of gall formation. While previous studies have recognized variation in gall morphology, most have attributed this primarily to the inducing insect species^21,22^. However, our findings align with Formiga *et al*. (2014), who suggested that stems exhibit lower morphoplasticity than leaves^25^. Building on their work, which examined one plant species, our study expands the comparative morphological analysis to 245 species across 26 plant orders, enabling broader phylogenetic validation for the influence of plant organ-derived factors on gall morphology. Our results suggest that the structural differences between stems and leaves impose inherent constraints on gall development, independent of the insect and plant species. Additionally, the use of fractal dimension analysis provides an objective and quantitative measure of morphological complexity, reducing biases associated with subjective descriptions. By leveraging images from iNaturalist, we were able to analyze a diverse and extensive dataset that would have been logistically challenging to compile through traditional fieldwork alone. Despite these strengths, our study has some limitations. First, the number of images used in our analysis (555) is relatively small compared to other iNaturalist studies (for example, the paper on ratsnake color patterns used over 8,000 images^6^). One reason for the small number is that both insect and plant species must be identified. In addition, while color patterns can be analyzed using RGB values, the analysis is limited to high-resolution images when considering the extraction of features with respect to morphology. In other words, the criteria for selecting images have become stricter in order to ensure that the morphological quantifications are reliable. Increasing the dataset size through improved data acquisition methods would enhance the robustness and increase the statistical power of future studies. Moreover, deep learning approaches have the potential to improve large-scale morphological analyses. Automated image recognition and classification could facilitate the processing of massive datasets, reducing manual screening efforts and increasing the consistency of morphological assessments^34^. Given the compatibility of deep learning with large, community-driven databases like iNaturalist, integrating machine learning models could revolutionize future studies on gall morphological diversity. Second, our dataset is biased toward well-documented regions such as North America and Europe, limiting the global applicability of our findings. Future studies should aim to incorporate data from underrepresented regions to obtain a more comprehensive understanding of gall morphology worldwide.

The reduced morphological complexity of stem-derived galls suggests that the physical and developmental characteristics of stems may limit the degree of morphological innovation achievable through insect-induced manipulation. Stems typically possess greater structural rigidity and lignification, especially with the secondary growth commonly observed in woody plants^35^, which may restrict cellular proliferation and reprogramming during gall induction. In contrast, leaves are developmentally more plastic, exhibiting higher cellular totipotency and greater sensitivity to hormonal and mechanical signals^36–38^, potentially allowing for more elaborate and variable gall morphologies. These differences in tissue plasticity may reflect fundamental developmental constraints that influence how plant organs respond to insect cues. From an evolutionary perspective, the greater morphological complexity observed in leaf-derived galls may have important implications for the diversification of gall-inducing insects^39^. Insects that exploit more developmentally flexible organs may be subject to fewer structural constraints, enabling a broader repertoire of gall forms and possibly enhancing ecological specialization or host manipulation. This view complements prior findings suggesting that leaf-derived galls tend to exhibit higher morphological innovation and host specificity than those induced on stems^21,25^. These results collectively highlight the importance of integrating both plant and insect factors, particularly plant tissue plasticity, when examining the evolutionary dynamics of gall formation and morphological diversity.

To move beyond correlative patterns, it is essential to understand the molecular mechanisms underlying organ-specific differences in gall morphology. Transcriptomic studies have revealed that gall-inducing insects broadly activate host developmental regulators involved in cell proliferation, meristem maintenance, and floral organ identity, suggesting a conserved “toolkit” used across taxa^22,41,42^. Hormone-related genes, particularly those in the auxin and cytokinin pathways, are also commonly upregulated ^43,44^. However, tissue-specific responses are evident: leaf-derived galls often suppress photosynthesis-related genes, while stem-derived galls downregulate cell wall biosynthetic pathways ^45,46^. These findings suggest that while gall-inducing insects initiate broadly similar developmental programs, the resulting morphologies depend on the structural and developmental context of the targeted plant organ. To test these hypotheses mechanistically, experimental systems are needed that allow manipulation of both plant and insect components. The Arabidopsis-based Gall-Forming Assay (Ab-GALFA) offers such a platform, enabling controlled exposure of model plants to insect-derived factors^47^. Applying this system to different plant tissues could help reveal whether the same insect signals elicit distinct transcriptional or morphological outcomes depending on plant organ.

In conclusion, our study indicates that plant organ shapes gall morphology regardless taxa. These findings highlight the importance of considering both plant and insect factors in understanding morphological diversity, and pave the way for mechanistic studies. This approach opens new avenues for investigating large-scale patterns of species interactions, functional morphology, and leveraging the power of publicly accessible datasets.

## Methods

### Data acquisition from iNaturalist and filtering steps

The workflow for data acquisition is illustrated in **Fig. S1**. Gall images and associated metadata were retrieved from iNaturalist.org (retrieval date: July 23, 2023) by selecting only observations labeled as “Research Grade” to ensure basic quality standards. Using the keyword “gall”, we initially obtained approximately 130,000 observations, and the metadata were downloaded via an R script using the *rinat* package^48^. To refine the dataset, duplicate insect species entries were removed, narrowing the dataset to 3,253 observations. Further filtering was applied to select only images that were of high resolution, captured from appropriate angles, and clearly showed the originating plant organ (leaf or stem), along with observations that included location metadata and taxonomic identification of both the host plant and gall-inducing insect at least at the order level. Given the variability in image quality and metadata completeness, manual screening was conducted to ensure that only images meeting these criteria were retained. After filtering, host plant species and insect species were identified to the family and order levels using taxonomic literature and expert assessment. The final dataset included 555 high-quality gall observations spanning 26 plant orders, and these data were used for subsequent morphological and phylogenetic analyses. All metadata for the final dataset, including taxonomic classifications, are provided in **Table S1**. To visualize the distribution of iNaturalist observation data, the map was created with the map_data()function in the *maps* package^49^ of R (v4.2.1) and added the latitude and longitude information coordinates with iNaturalist observation.

### Data acquisition from a gall book and filtering steps

Gall images and metadata were also collected from “Plant Galls of the Western United States” by Ronald A. Russo^27^. Non-insect-induced galls, such as those caused by bacteria, fungi, nematodes, and mistletoe, were excluded, along with galls of unknown inducers. Ultimately, 422 out of 536 entries were retained for analysis, with exclusions primarily due to missing inducer information, non-insect gall inducers, or insufficient data on host plant identification. Each gall was annotated with the order name of its host plant and gall-inducing insect, as well as the originating plant organ. Metadata for the gall book data are listed in **Table S2**.

### Comparison of host plant and gall-inducing insect proportions between data sources

The order-level proportions of insect and plant species were extracted from both datasets, and their total percentages were calculated. Correlation analyses were conducted for the number of insect and plant orders in each data source, using both raw counts and log10-transformed values. Pearson’s correlation coefficients were calculated using R (v4.2.1).

### Lineage distribution of the data retrieved from iNaturalist

To assess which insect-plant combinations were more common in the iNaturalist data obtained, the number of insect-plant combinations at order level was counted and indicated as a histogram. Heatmap also indicated the number of insect-plant combinations in a log scale with removing single combinations of insect-plant.

### Fractal dimension analysis for quantifying gall morphological complexity

Gall images were manually classified as leaf-derived or stem-derived ones. Each gall shape was binarized using Photoshop (v26.4.1). Morphological complexity was quantified using fractal dimension analysis with the box-counting method applied to binarized images^50,51^. We imported JPG files with the import_jpg() function in the *Momocs* package^52^ and used the fd.estim.boxcount() function to estimate the fractal dimension in the *fractaldim* package^53^. The analyses were conducted using R (v4.3.2).

### Principal component analysis of gall morphological traits

Principal component analysis (PCA) was conducted using four quantitative traits and two categorical traits from 50 randomly selected gall images. The measured quantitative traits included: (1) aspect ratio, (2) relative gall thickness compared to the originating plant organ, (3) similarity to an ellipse, and (4) number of spines. The categorical traits included (5) color difference between the gall and the originating plant organ, and (6) density and length of trichomes. For color difference, a value of 1 was assigned when the gall exhibited a color change compared to the originating organ (leaf or stem), and 0 when no color change was observed. For trichome density and length, a categorical scale was used: 0 for no trichomes, 1 for sparse and short trichomes, 2 for sparse and long trichomes, 3 for dense and short trichomes, and 4 for dense and long trichomes. ImageJ (v1.51) was used for trait measurement. PCA was performed using the prcomp() function in R (v4.2.1) with variables centered and scaled to unit variance (`scale = TRUE`). Raw data of all the measured traits are listed in **Table S4**.

### Data analysis and statistical analysis

The following analyses and statistical tests were conducted in R (v4.2.1) using the functions of cor() and wilcox.exact(). Wilcoxon rank sum tests with Bonferroni correction were used for multiple comparisons. To quantify the effect of host plant organ on gall morphological complexity while accounting for phylogenetic relatedness, we used a Bayesian generalized linear mixed model implemented in the R package *MCMCglmm*^54^. The response variable was the fractal dimension value of stem-, and leaf-derived gall, and the explanatory fixed effect was plant organ (leaf or stem), treated as a categorical variable. To control for potential phylogenetic non-independence, we included the phylogenetic relationships of gall-inducing insect orders and host plant orders as random effects. The insect phylogeny was incorporated using a phylogenetic covariance matrix derived from an ultrametric tree at the order level, specified via the `ginverse` argument. For host plants, taxonomic order was included as a random effect without a specified phylogenetic structure, thereby modeling inter-order variation as independent and identically distributed. All models assumed a Gaussian distribution for the response variable and were run for 13,000 iterations with a burn-in of 3,000 and a thinning interval of 10. Priors for random effects were specified as weakly informative inverse-Wishart distributions (*V = 1*, ν*= 0*.*002*). Posterior distributions were summarized to obtain the posterior mean, 95% credible intervals, and the MCMC-derived *p*-value (p_MCMC_) for fixed effects. Significance was inferred when the 95% credible interval did not include zero and/or when p_MCMC_ < 0.0*5*. Models were compared using the deviance information criterion to assess the relative contribution of plant organ identity and phylogenetic structure to morphological variation.

## Acknowledgments

We thank Mr. Shunsuke Uda, Ms. Jyothi Naga Udandarao, and Ms. Mina Atarashi for helping to summarize the data from the book of plant galls. We appreciate Prof. Keiko Torii for sharing the gall book for our research.

## Data availability statement

All data are provided in the supplemental file.

## Author Information

These authors contributed equally: Kanako Bessho-Uehara and Riki Takara

## Author contributions

KBU, RT, and KT designed this study and wrote the manuscript. RT, KS, and KBU retrieved and evaluated data from iNaturalist and books. KBU, RT and KT performed image analysis and statistical analysis. All authors have reviewed and commented on the manuscript.

## Corresponding author

Correspondence to Kanako Bessho-Uehara.

